# Divergent evolution of insecticidal protein families across bacteria and ferns

**DOI:** 10.1101/2025.11.17.688980

**Authors:** Haruka Endo, Hokto Kazama

## Abstract

Insecticidal Cry proteins produced by the bacterial insect pathogen *Bacillus thuringiensis* possess conserved three domains (3D). Two other families of insecticidal proteins—Prb proteins from bacterial symbionts of entomopathogenic nematodes and a marine bacterial pathogen, and IPD113 proteins from ferns—are two-domain (2D) proteins structurally resembling domains I and II of Cry proteins. Despite the structural resemblance, the amino acid sequence identity among these families is only ∼10%, supporting the prevailing view that they evolved independently through convergent evolution. However, the possibility of remote homology and ancient horizontal gene transfer (HGT) has remained largely unexplored. Here, we present evidence supporting divergent evolution of these protein families. We found that proteins containing Cry-like two domains are distributed far more broadly across bacteria, eukaryotes, and archaea than previously recognized. Phylogenetic analyses separated the structurally similar proteins into five clades including Cry, IPD113, Prb, and 2D/3D-mixed clades. Domain-level trees revealed intermixing within domain II, with nematocidal Cry proteins frequently clustering with IPD113 and 2D/3D-mixed clades, suggesting domain swapping events across these clades. Conserved amino acid motifs and domain core structures across clades further support a shared evolutionary origin. Phylogenetic and structural analyses suggest a bacterial origin of the fern IPD113 gene and its horizontal transfer via fungal symbionts, as well as multiple HGT events into diverse terrestrial and marine eukaryotes and archaea. This work redefines the evolutionary landscape of the insecticidal protein families, revealing their unexpectedly wide taxonomic distribution, possibly shaped through bacterial ecological networks and close biological interactions such as symbiosis.

## Introduction

The bacterial pathogen *Bacillus thuringiensis* (Bt) produces insecticidal and nematocidal Cry proteins, which have been widely used in agriculture for pest control for a century. The Cry protein family is highly diverse, consisting of 53 subfamilies that share less than 45% amino acid identity with each other (Crickmore et al. 2021). These proteins kill their specific targets including lepidopteran caterpillars, mosquitoes, beetles, and nematodes (van Frankenhuyzen 2009). Cry proteins are α-pore-forming toxins that share three conserved domains in their active form (Fig. 1A) (Grochulski et al. 1995). Domain I is a pore-forming domain composed of seven α-helices. Domain II adopts a β-prism structure that recognizes target membrane proteins or glycolipids in the host gut and plays a major role in determining host specificity. Domain III is a β-sandwich structure thought to contribute to structural stability and host recognition, although its precise role remains unclear.

**Fig 1.**
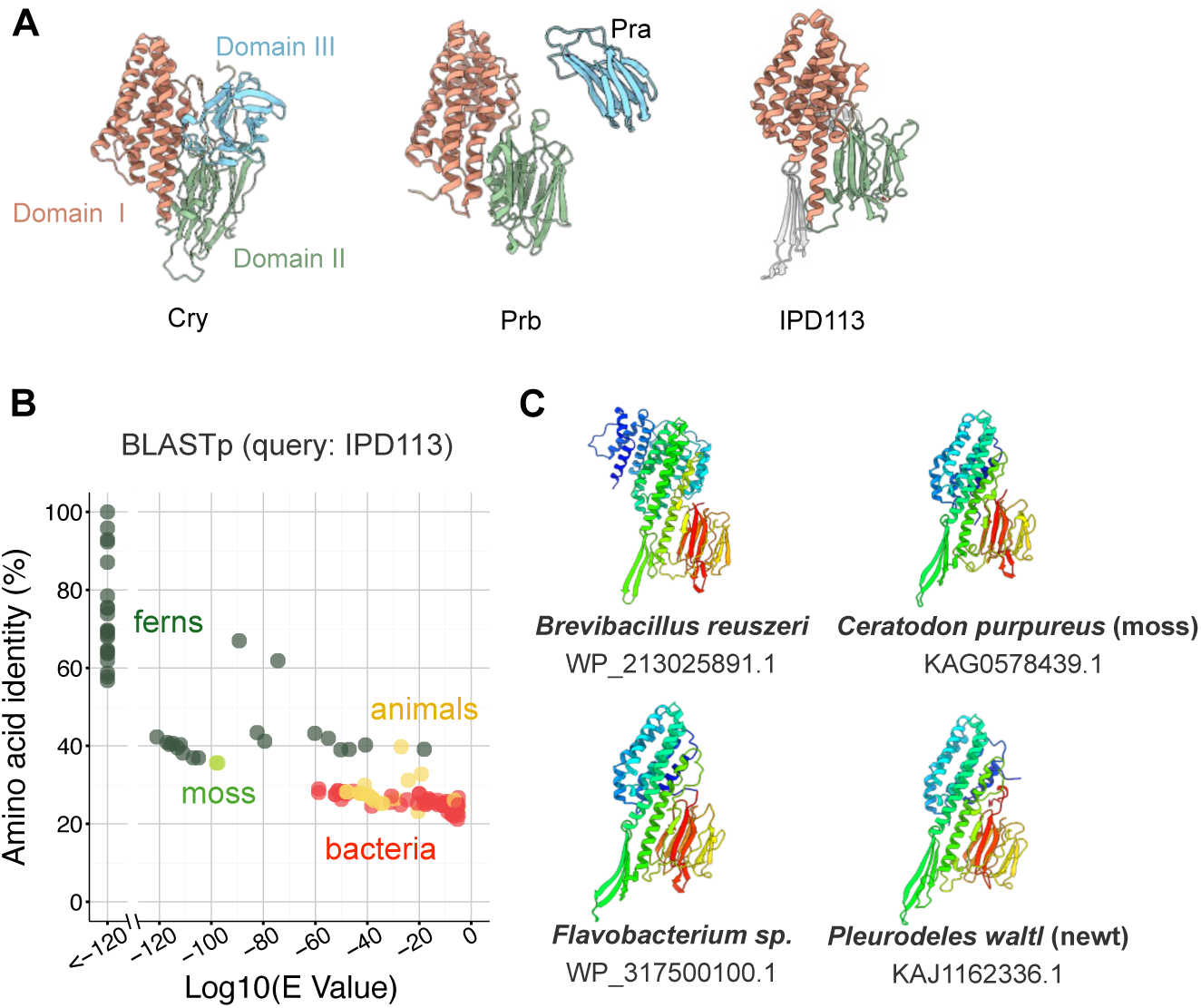
Putative IPD113 homologues identified by BLASTp. (A) Domain architecture of Cry, Prb, and IPD113 proteins. Cry (PDB: 1CIY), Prb (PDB: 7FDP), Pra (PDB: 7FCN), and IPD113 (PDB: 8D2J). (B) Scatter plots of amino acid identity versus E-value from BLASTp searches. *Leptochilus wrightii* IPD113 protein. Dots indicate putative homologues (E-value < 1e−5). Raw BLASTp results are provided in Table S1. (C) AF3-predicted structures of putative IPD113 homologues.

Interestingly, two other insecticidal protein families, Prb and IPD113, exhibit striking structural similarity to domains I and II of Cry proteins (Fig. 1A) (Wei et al. 2023; Prashar et al. 2023; Lee et al. 2015). Prb proteins (*Photorhabdus* insect-related toxin B component) are produced by bacterial symbionts of entomopathogenic nematodes, such as *Photorhabdus* and *Xenorhabdus*, as well as by the marine pathogen *Vibrio parahaemolyticus* (Waterfield et al. 2005; Duchaud et al. 2003; Lee et al. 2015). Prb proteins couple with another component Pra to exhibit toxicity toward lepidopteran caterpillars (*Photorhabdus* and *Xenorhabdus*) and shrimp (*Vibrio*). In contrast, IPD113 proteins are found in polypodiopsida ferns and are toxic to lepidopteran caterpillars as single-component toxins, without the need for additional protein partners (Wei et al. 2023). Despite their structural resemblance, Cry, Prb, and IPD113 proteins share only ∼10% amino acid identity. Such low amino acid identity is often referred to as the “midnight zone”, where most protein pairs are unrelated in both structure and function (Rost 1997). In this zone, it is generally difficult to determine whether proteins have originated from a common ancestor through divergent evolution, or independently acquired similar structures through convergent evolution, or whether the similarity is merely coincidental. Given the low sequence identity and the presence of introns in IPD113 genes, the original report on IPD113 favored the hypothesis that these proteins represent a case of convergent evolution (Wei et al. 2023). However, the possibility of remote homology and ancient horizontal gene transfer (HGT) has remained largely unexplored.

In this study, we aimed to examine whether the three insecticidal protein families (Cry, Prb, and IPD113) are evolutionarily linked. By aligning AlphaFold 3 (AF3)-predicted structures, we were able to accurately extract amino acid residues corresponding to the two domains even among proteins with very low amino acid identity. Although the extracted sequences remain highly divergent, pairwise alignment-based phylogenetic analysis developed for protein superfamilies enabled reliable tree construction. The results indicate that Cry, Prb, and IPD113 proteins share a common evolutionary origin, and that their genes have been unexpectedly disseminated across diverse bacterial, archaeal, and eukaryotic lineages via HGTs, some of which may have been mediated by close ecological interactions such as symbiosis.

## Results

### Proteins containing Cry-like two domains are broadly distributed across taxa

To explore the possibility of remote homology, we performed BLASTp searches using IPD113 and Prb sequences as queries. Putative IPD113 homologues (E-value < 1e-5) were identified in ferns, mosses, bacteria, and animals, including frogs and newts (Fig. 1B and Table S1). These bacterial and animal proteins share approximately 30% amino acid identity with the query sequence (GenBank: UDN67319.1 from *Leptochilus wrightii*) (Fig. 1B), and their AF3-predicted structures resemble those of fern IPD113 proteins (Fig. 1C). A subsequent BLASTp search using a bacterial IPD113-like protein (GenBank: WP_119147942.1 from *Cohnella faecalis*) as the query revealed additional IPD113-like proteins in fungi, archaea, and protists, as well as in plants, bacteria, and animals (Fig. S1A and Table S1). Similarly, BLASTp searches with Prb sequences from *Photorhabdus akhurstii* and *Vibrio parahaemolyticus* (GenBank: AEQ33638.1 and 3X0U_A) identified putative homologues in both bacteria and insects, such as beetles and aphids (Fig. S1B and Table S1). Eukaryotic genes encoding these proteins often possess introns (Table S2), suggesting that they are not the result of bacterial contamination in sequencing but are integrated and functional components of the eukaryotic genome, similar to fern IPD113 proteins. Together, these findings suggest an unexpectedly broad taxonomic distribution of proteins containing Cry-like two domains.

### Phylogenetic analysis identifies the 2D/3D-mixed clade

To reconstruct the phylogenetic relationships among proteins containing two Cry-like domains in the midnight zone, we predicted their structures using AlphaFold3 (Abramson et al. 2024), aligned them with FoldMason (Gilchrist, Mirdita, and Steinegger 2024), and extracted the amino acid residues corresponding to domains I and II (see Methods). We selected 557 non-redundant sequences with ≥95% identity that fully contain the two domains. These include 268 Cry proteins with quaternary rank 1, i.e., the canonical isoform denoted by the terminal “1” in Cry names (e.g., Cry1Aa<u>1</u>, Cry3Bb<u>1</u>), listed in the Bacterial Pesticidal Protein Resource Center (www.bpprc.org), as well as six sequences identified by BLASTp using Cry1Aa1 (GenBank: AAA22353) as a query.

Since the extracted sequences are highly divergent, conventional multiple sequence alignment (MSA)-based phylogenetic methods, such as maximum likelihood, were not applicable. Instead, we employed the Graph Splitting (GS) method, a graph-based approach suitable for reconstructing superfamily-scale phylogenetic trees (Matsui and Iwasaki 2020). The resulting GS tree grouped the proteins containing the Cry-like two domains into five clades (Fig. 2A). Based on known protein families, we designated three of these clades as Cry, IPD113, and Prb, and the remaining two as the 2D/3D-mixed and *Photobacterium* clades (Fig. 2A). The Cry clade comprises only 3D Cry proteins and includes the majority of known Cry proteins (Fig. 2B). In contrast, the 2D/3D-mixed clade contains both the remaining 3D Cry proteins (Cry2, Cry11, Cry18, Cry63, and some members of Cry5 and Cry21) and proteins from diverse non-Bt bacteria including Pseudomonadota, as well as from fungi and archaea (Fig. 2A and Fig. S2A). While most non-Bt members of this clade are Cry-like two domain proteins, some harbor one or two additional domains (Fig. 2B). The IPD113 clade includes members from non-Bt bacteria, fungi, plants, animals including frogs and newts, archaea, and protists (Fig. 2B and Fig. S2B). Some members of this clade possess additional domains beyond the canonical two. The Prb clade includes 2D proteins from *Photorhabdus* and *Xenorhabdus* as well as from Pseudomonadota (e.g., *Serratia*, *Yersinia*), along with 3D proteins from beetles and aphids (Fig. S2C). These insect-derived 3D proteins have a Pra-like N-terminal domain followed by two Prb-like domains (Fig. S1C). The Photobacterium clade contains only four 2D proteins, all from *Photobacterium* species, while another *Photobacterium*-derived sequence belongs to the 2D/3D-mixed clade (Fig. 2B). These results suggest an evolutionary link between 3D Cry proteins and the 2D proteins in the 2D/3D-mixed clade. The phylogeny also indicates that the Prb and IPD113 families are closely related.

**Fig 2.**
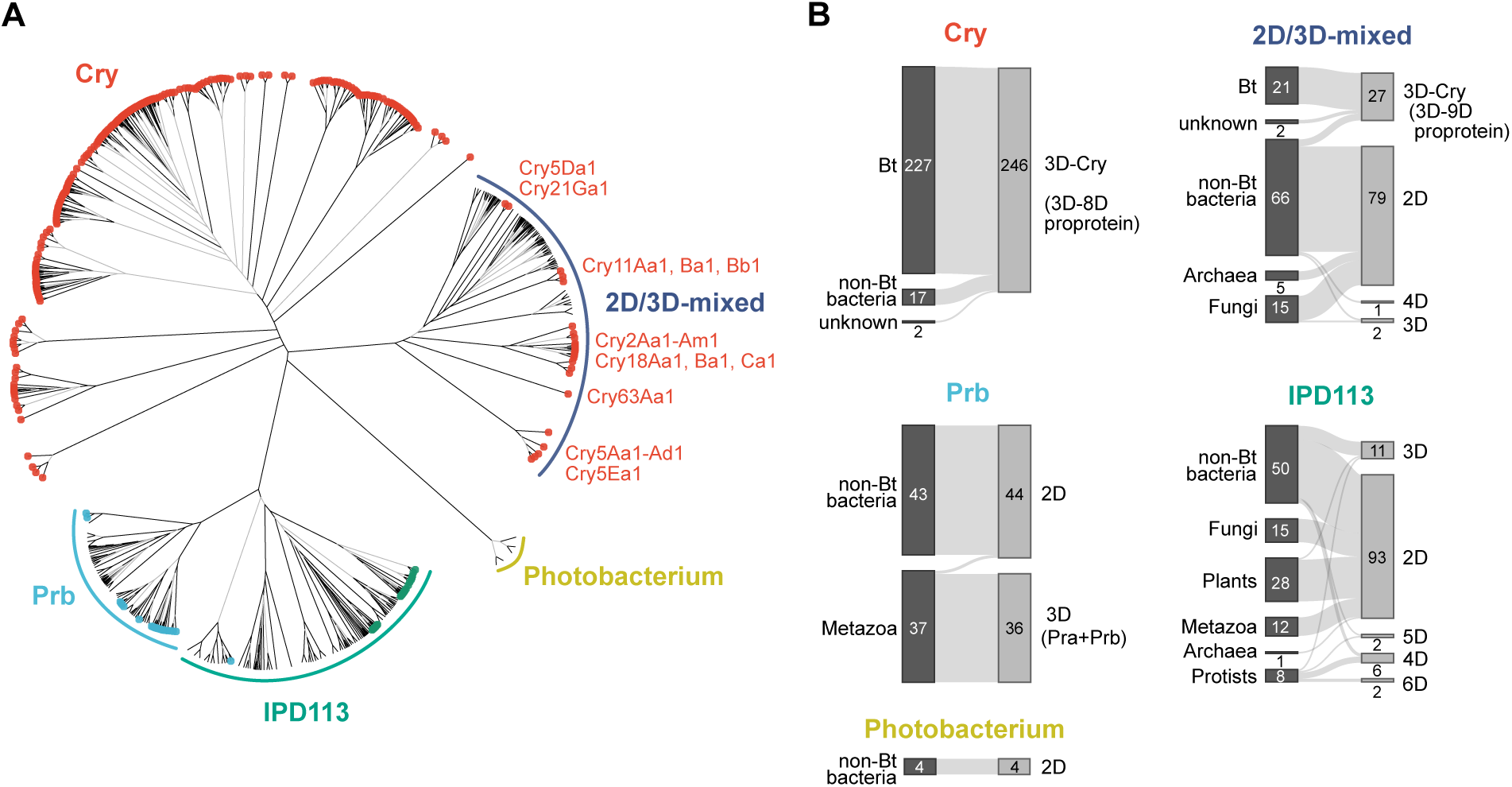
The GS tree of domain I–II region of proteins containing Cry-like two domains. (A) Graph Splitting tree of the domain I proteins containing Cry-like two domains. Black and gray branches indicate high and low reliability, supported by edge perturbation (EP) values ≥ 0.9 and < 0.9, respectively. Red, light blue, and green node circles indicate known Cry proteins, Prb proteins from *Photorhabdus*, *Xenorhabdus*, and *Vibrio*, and fern IPD113 proteins. (B) Sankey plots showing the taxonomy and the number of domains, based on AF3-predicted structures, in each clade.

### Domain II swapping across clades except for Prb

We next generated domain-level phylogenetic trees. While the domain I tree largely recapitulated the topology of the two-domain tree (Fig. 3A), the domain II tree revealed extensive intermixing among Cry, IPD113, 2D/3D-mixed, and Photobacterium clades (Fig. 3B), suggesting that domain II swapping events occurred across these four clades. Specifically, several nematocidal Cry proteins (Cry5A, Cry5D, Cry5E, and Cry12), as well as Cry31 and Cry70, clustered far from other Cry proteins and instead grouped with members of other clades, such as IPD113. Since domain swapping typically requires homologous recombination, it is likely that the genes encoding these proteins contained complementary sequences flanking the domain II coding region. These findings support the possibility that proteins with Cry-like domains are remote homologues.

**Fig 3.**
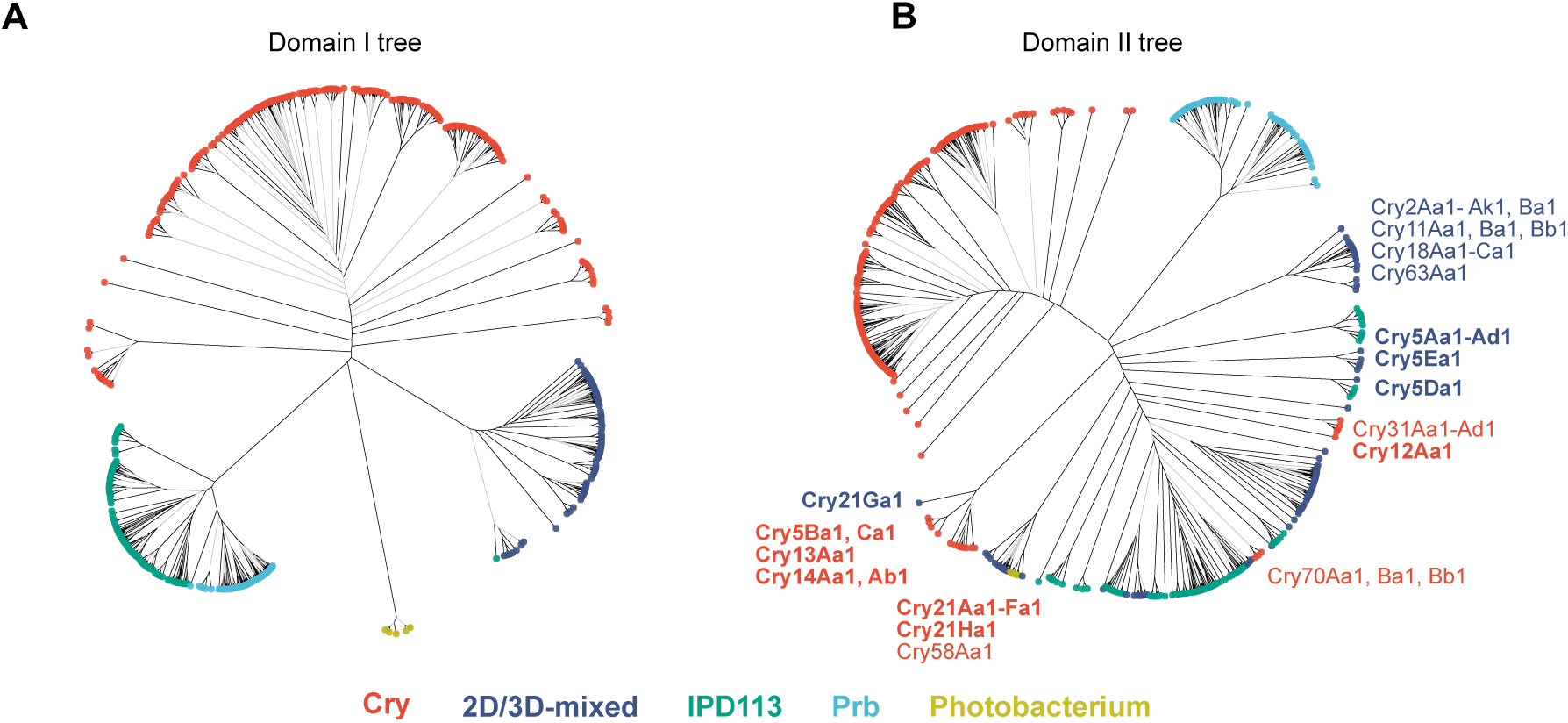
The domain-level GS tree of proteins containing Cry-like two domains. GS trees of domain I (A) and domain II (B) sequences. Black and gray branches indicate high and low reliability, supported by edge perturbation (EP) values ≥ 0.9 and < 0.9, respectively. Node circle colors correspond to the clades defined in the two-domain GS tree shown in Fig. 1C. Cry proteins shown in bold indicate nematocidal Cry proteins.

### Proteins containing Cry-like two domains share amino acid motifs across clades

To examine remote homology at a resolution beyond the domain level, we performed a search for conserved amino acid motifs. Among the 60 motifs identified by MEME (Bailey et al. 2015) (Fig. S3A), 26 motifs were shared across at least two clades (Fig. 4A). They locate within domain I α-helices, the linker region between domains I and II, and domain II β-strands, with their relative positions being largely conserved (Fig. S4A). t-SNE visualization of motif patterns revealed partial intermixing between clades in all cases, whether motifs were derived from domain I alone, domain II alone, or the domain I–II region (Fig. 4B–D).

**Fig 4.**
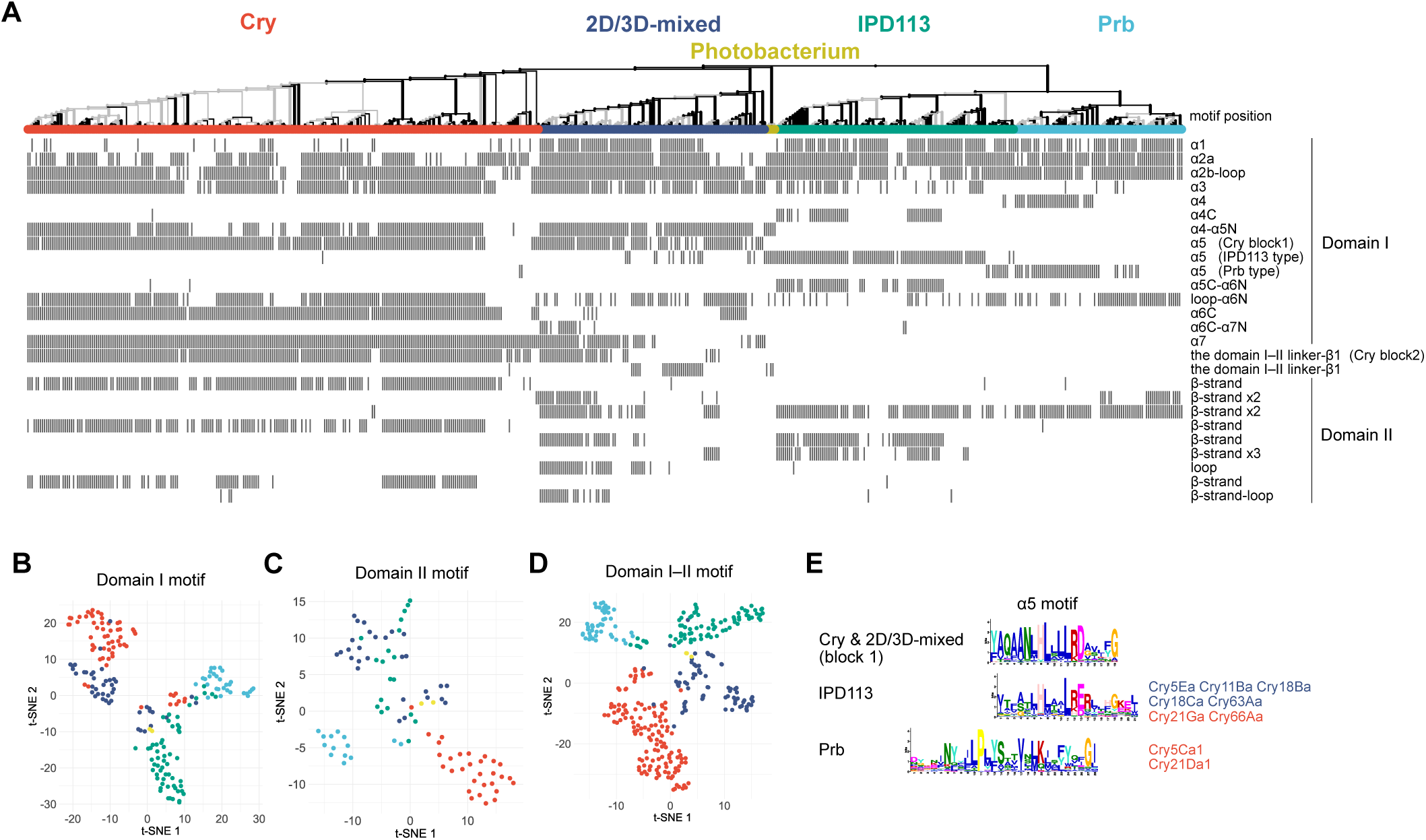
Amino acid motif patterns. (A) Raster plot of amino acid motif patterns. Twenty-six motifs shared across at least two clades were mapped onto the two-domain GS tree shown in Fig. 2A. (B–D) t-SNE visualization of motif patterns. The motif presence/absence matrix from (A) was dimensionally reduced to visualize motifs located in domain I (B), domain II (C), and the domain I–II region (D). (E) α5 motifs of each clade. Cry proteins carrying IPD113-type or Prb-type α5 motifs are listed.

On average, the proteins in each clade possess more than four shared domain I motifs, each approximately 20 amino acids in length (Fig. S4B), suggesting that domain I is unlikely to have been acquired through convergent evolution. In line with this, the α5 motif, located at the center of the domain I α-helical bundle, shows a gradual transition across clades (Fig. 4A and E). In Cry proteins, the α5 motif corresponds to block 1, one of the five conserved sequence blocks (Höfte and Whiteley 1989). 79% of the 2D/3D-mixed clade members including non-Cry proteins retain the Cry-type α5 motif. Furthermore, 50% and 37% of these members also share the α7 and the domain I–II linker–β1 motifs, respectively, both of which correspond to Cry block 2 (Fig. 4A and E). Most members of the IPD113 clade possess an α5 motif that is similar, though not identical, to the Cry-type α5 motif, while some Cry proteins in both the Cry and 2D/3D-mixed clades carry the IPD113-type α5 motif. Members of the Prb clade, on the other hand, have a longer α5 motif that is distinct from both the Cry- and IPD113-types. Notably, Cry5Ca1 and Cry21Da1 classified in the Cry clade as well as some members of the IPD113 clade, contain the Prb-type α5 motif. Since the motifs flanking the IPD113-type α5 motif in Cry5Ea1 and Cry21Ga1 and the Prb-type α5 motif in Cry5Ca1 and Cry21Da1 are similar to those of closely related Cry5 and Cry21 proteins (Fig. S4C), it is plausible that either a subdomain-level swapping event occurred, or the surrounding motifs emerged after acquiring the IPD113-type and Prb-type α5 motif. This observation further suggests an evolutionary relationship of domain I across Cry, IPD113 and Prb proteins.

In contrast, shared motifs are less prevalent in domain II, with 20–40% of sequences in the IPD113, 2D/3D-mixed, and Photobacterium clades lacking any such motifs. Although only Prb proteins showed no signatures of domain swapping in domain II (Fig. 3B), they nevertheless possess two long conserved motifs, each corresponding to a pair of β-strands—one shared with non-Cry 2D/3D-mixed proteins and the other shared with both non-Cry 2D/3D-mixed and IPD113 proteins (Fig. 4A). These findings suggest a common ancestry of domain II across these clades.

### Domain core structures are highly conserved beyond clades

To further investigate signatures of divergent evolution, we performed structural analyses, as protein structures are more conserved than amino acid sequences (Illergård, Ardell, and Elofsson 2009). For structural analyses, we selected 498 predicted models with high confidence, defined as an average pLDDT ≥ 80 and PAE ≤ 10 Å in both domains I and II. We compared the domains I and II of *P. akhurstii* Prb and *L. wrightii* IPD113 with those of other predicted structures. Although sequences from other clades exhibited low sequence scores, some of them still fell within the root-mean-square deviation (RMSD) range observed for comparisons within the Prb or IPD113 clade (Fig. 5A-D). When aligned pairwise, we found that even among structures from other clades, approximately 25–50% of the Cα atoms in each domain could be superimposed within 1.5 Å (Fig. 5E–H). In domain I, the central helix α5 together with several adjacent helices showed strong structural conservation (Fig. 5I and J). In domain II, one or more of the antiparallel β-sheets forming the β-prism overlapped almost completely (Fig. 5K and L). The domain-level GS tree based on the 3D interaction (3Di) alphabet, a type of structural alphabet that describes tertiary interactions in FoldMason (van Kempen et al. 2023), revealed partial intermixing among clades (Fig. S5). These cross-clade conservations of domain core structures support the notion that these domains have undergone divergent evolution while maintaining their functional architecture, rather than having arisen through convergent evolution.

**Fig 5.**
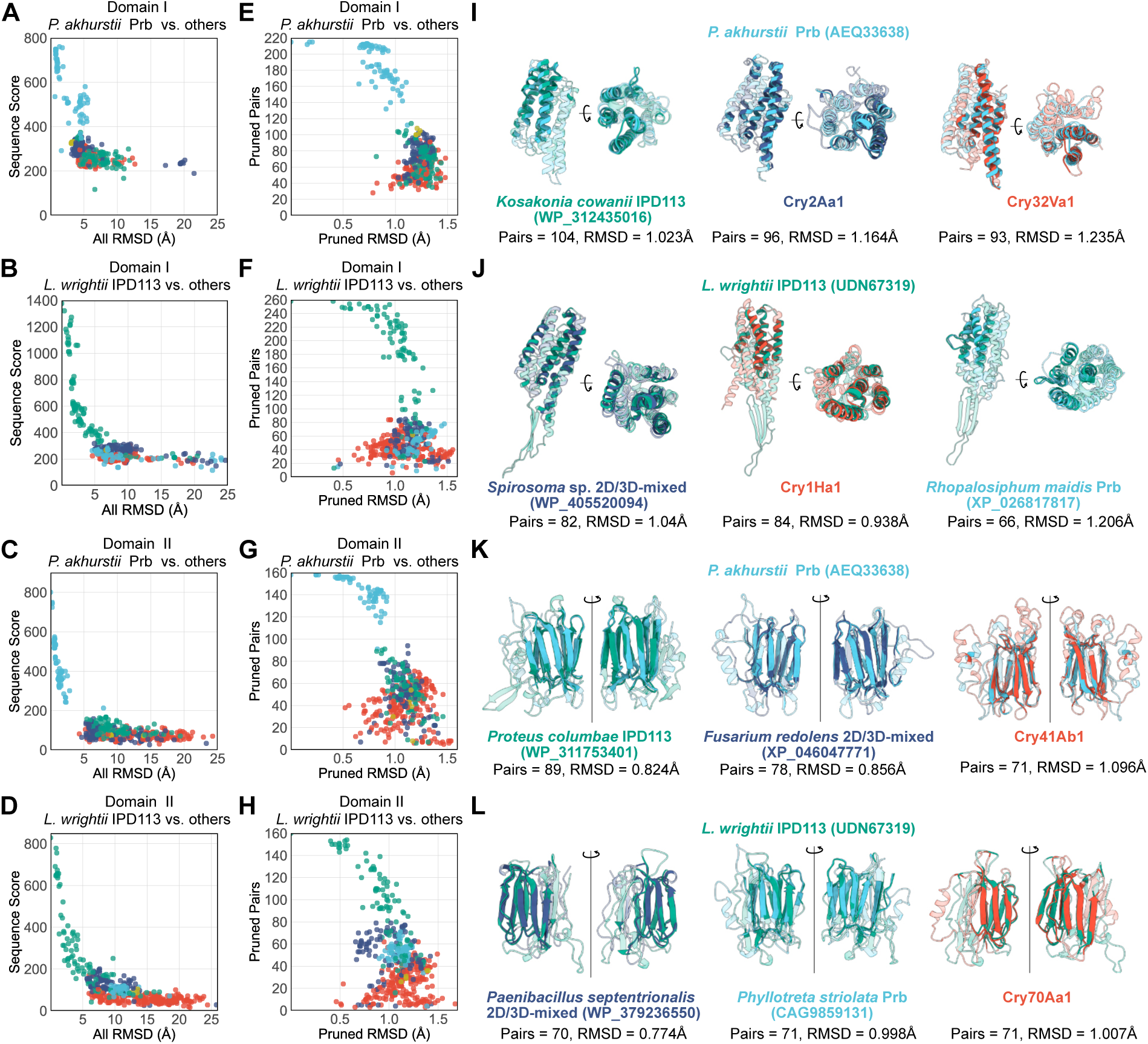
Domain-level structural comparisons. The domain I and II structures of *Photorhabdus akhurstii* Prb and *Leptochilus wrightii* IPD113 were compared with those of other proteins containing Cry-like two domains. (A–D) Sequence similarity scores and whole-domain RMSD values obtained from pairwise alignments and structural superpositions. Dot colors correspond to the same clade color scheme shown in Fig. 1C. (E–H) Pruned residue pairs and their pruned RMSD values derived from structural superpositions. Dot colors correspond to the same clade color scheme shown in Fig. 1C. (I–L) Representative structural superpositions with proteins from different clades. Pruned regions are highlighted, and non-pruned regions are shown with increased transparency.

### Cry domain III and Pra are not evolutionarily related

To test the homology among Cry domain III, Pra, and the Pra-like domains of insect Prb-containing proteins, we performed the same analyses on these domains. The GS tree based on amino acid sequences separated Cry domain III from Pra and the insect Pra-like domains, while Pra and the insect Pra-like domains formed a single clade (Fig. S6A). Motif analysis revealed that Cry domain III did not share motifs with Pra or the insect Pra-like domains (Figs. S6C and D). Meanwhile, Pra and the insect Pra-like domains shared two motifs (Fig. S6C and D). The GS tree based on structure (3Di) also showed intermixing between Pra and the insect Pra-like domains, but not with Cry (Fig. S6B). In pairwise structural alignments with *P. akhurstii* Pra, some Cry domain III showed pruning of approximately 30% of the domain length; however, no conservation of the core structures observed in domain I or II was detected (Fig. S6E-G). These results suggest that Cry domain III and Pra are not remote homologues, whereas Pra and the insect Pra-like domains share a common evolutionary origin. *Pra* and *Prb* genes are located adjacent to each other in the *Photorhabdus* genome (Waterfield et al. 2005), suggesting that the two proteins have fused into a single protein at some point.

### Many horizontal gene transfers shaped broad taxonomic distribution

As our results suggested a common evolutionary origin of Cry, IPD113, and Prb, we hypothesized that ferns, as well as other eukaryotes and archaea, may have acquired these genes through ancient HGT. In the GS tree shown in Fig. 1C, the 2D/3D-mixed and IPD113 clades include a mixture of bacterial and non-bacterial clades, and the Prb clade contains a single bacterial member nested within an animal clade (Fig. S7). Since the GS method does not estimate evolutionary distances, we additionally constructed maximum-likelihood (ML) trees for each clade to obtain amino acid distances under ML substitution models. In this analysis, we used only the sequences that were subjected to structural comparisons. As expected, approximately 30–40% of all branches in ML trees lacked strong support (Fig. 6A and S9). To narrow down the possible scenarios of ancient HGT, we compared amino acid (AA) distances with structural distances (Fig. S8). We found that a number of pairs between bacteria and non-bacterial taxa exhibit a lower 3Di/AA distance ratio than the regression curves derived from bacteria–bacteria pairs within the same clade. This indicates that, for the same amino acid distance, structures of those pairs are closer than expected for typical bacterial pairs. Such structural closeness supports the placement of non-bacterial members within bacterial clades and thus suggests ancient HGT events. In particular, pairs that also show shorter AA distances (below the bacterial mean) can be regarded as stronger candidates, because lower sequence divergence combined with high structural similarity between bacteria and non-bacterial taxa is more parsimoniously explained by relatively recent HGT than by independent evolution.

**Fig 6.**
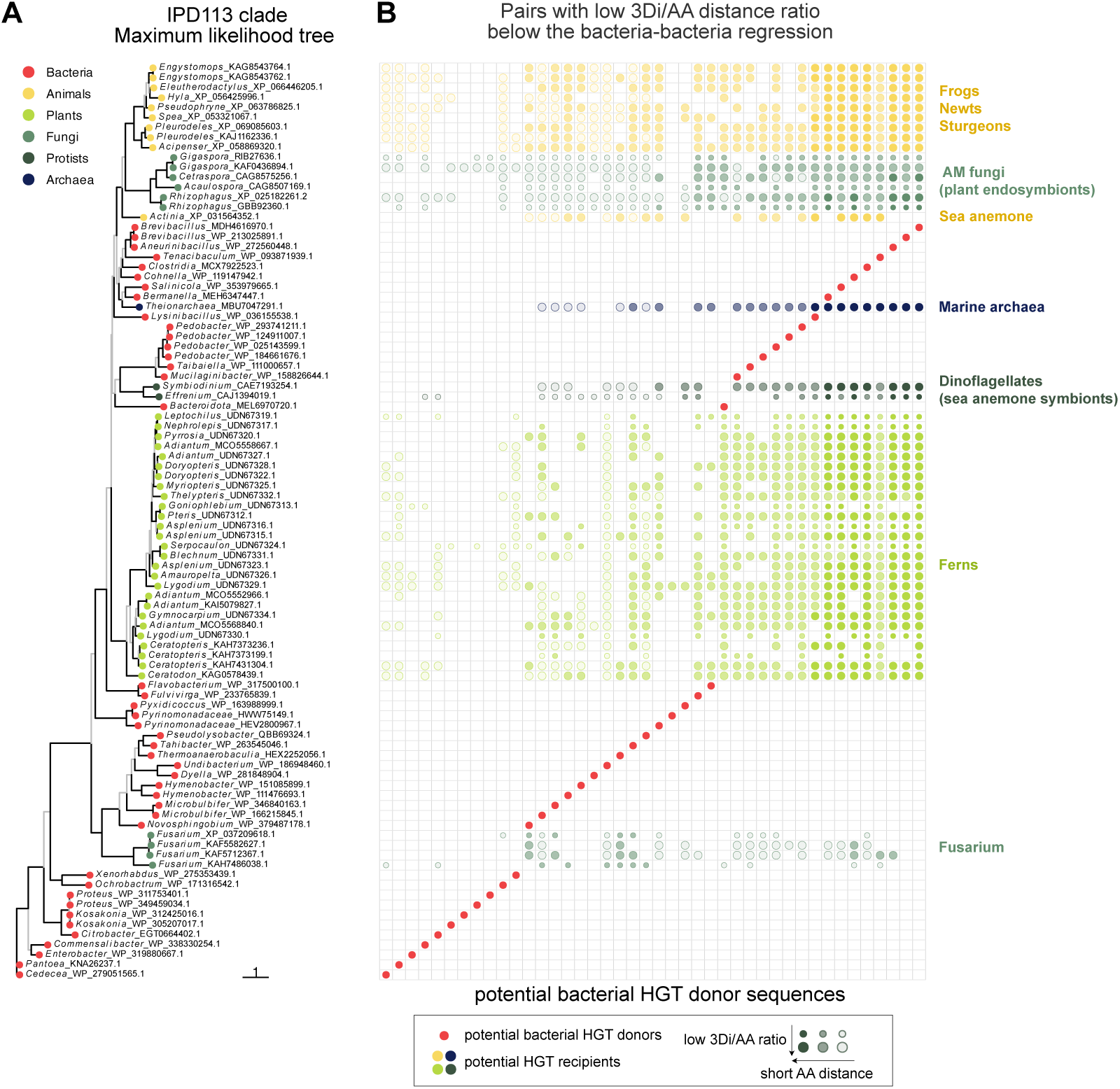
Potential HGT pairs in the IPD113 clade. (A) Maximum likelihood tree of the IPD113 clade. Black branches indicate high reliability (Ultrafast Bootstrap ≥ 95% and SH-aLRT ≥ 80%); other branches are shown in gray. Node circle colors indicate taxonomy. The scale bar represents the number of substitutions per site. (B) Pairs with a low 3Di/AA distance ratio, falling below the bacteria–bacteria regression. The vertical position of each circle corresponds to the sequence labels in the phylogenetic tree shown in (A). Each column corresponds to a red-highlighted bacterial sequence (potential HGT donor), with which sequences of non-bacterial sequences (rows) are compared. Proteins plotted in the same column have a 3Di/AA distance ratio that falls below the bacteria–bacteria regression in Fig. S8, indicating potential HGT recipients. The fill opacity of each circle reflects its amino acid (AA) distance relative to the mean of bacteria–bacteria pairs: circles with AA distances above the mean are outlined with high transparency, those between the mean and −1 SD are semi-transparent, and those below −1 SD are shown as fully opaque. Large circles indicate proteins below −1 SD from the regression, while small circles indicate those between 0 and −1 SD. In particular, large opaque circles are more likely to represent candidate HGT pairs involving bacterial sequences.

In the IPD113 clade, we found bacteria–fern pairs that met the criteria (Fig. 6), suggesting a bacterial origin for IPD113. Specifically, seven bacterial sequences from *Brevibacillus*, *Aneurinibacillus*, *Clostridia*, *Cohnella*, *Salinicola*, and *Bermanella*, showed a low 3Di/AA distance ratio and a short AA distance (both below the bacterial mean minus one standard deviation) (Fig. 5B, large solid yellow-green circles), making them a strong candidate HGT donor. Interestingly, most of the bacteria that emerged as potential HGT donors to ferns also exhibited a lower 3Di/AA distance ratio with arbuscular mycorrhizal (AM) fungi, known as plant endosymbionts, implying a scenario of bacteria–AM fungi–fern HGT. Likewise, bacteria–symbiont–sea anemone HGTs were suggested, as well as HGTs to frogs, newts, sturgeons, and *Fusarium* (Fig. 6). In the 2D/3D-mixed clade, additional HGT events were suggested toward *Fusarium* and other fungal plant pathogens and parasites, nematode-trapping fungi, and marine archaea (Figs. S9A–B). In the Prb clade, HGTs from bacteria, including the insect endosymbiont *Rickettsia*, to aphids and beetles were also suggested (Figs. S9C–D). However, compared with the distribution among bacteria–bacteria pairs, the bacteria–animal pairs tended to show larger AA and 3Di distances in this clade (Fig. S8C and F), suggesting that the actual HGT donors may include yet-unidentified bacterial lineages harboring Pra+Prb three-domain proteins. These results suggest that, in addition to HGT among diverse bacteria, close ecological interactions such as symbiosis have contributed to shaping the current broad taxonomic distribution of these genes.

## Discussion

Since Cry proteins have so far been identified only in *B. thuringiensis* and several other Bacillota bacteria including *Paenibacillus* and *Lysinibacillus*, they have generally been considered to represent a lineage uniquely acquired and diversified within these bacteria. Here, however, we show that the first two domains are broadly distributed across diverse organisms, with the Cry protein family representing one terminal outcome of this broader diversification. Although our results cannot determine whether the three-domain Cry proteins evolved by gaining an additional domain from two-domain ancestors or two-domain proteins emerged by domain loss from three-domain ancestors, the phylogenetic and motif analyses (Figs. 2–4) suggest that the 2D/3D-mixed clade has played an intermediate, bridging role linking the Cry, IPD113, and Prb lineages. In the 2D/3D-mixed clade, both 2D proteins and canonical 3D Cry proteins were found in *Paenibacillus* species (Table S2), suggesting that this clade represents a site of domain-3 gain or loss. In addition, certain *Lysinibacillus* species produce 2D proteins that belong to the IPD113 clade (Table S2).

Domain swapping has been proposed to underlie the diversification of Cry proteins (de Maagd, Bravo, and Crickmore 2001). Our results extend this concept beyond the Cry family, encompassing proteins in the 2D/3D-mixed, IPD113, and Photobacterium clades as well. Many Cry proteins that form intermixed clades are nematocidal (Fig. 3B), suggesting that ecological interactions may have occurred across bacteria including Bt strains that use nematodes as hosts. In contrast, domain II swapping appears not to have occurred with the Prb clade. This may be explained by the fact that *Prb* genes are chromosomally encoded in *Photorhabdus*, *Xenorhabdus*, *Sodalis*, and *Yersinia* (Wang et al. 2023) whereas Cry proteins are typically plasmid-borne. Furthermore, *Photorhabdus* and *Xenorhabdus* are endosymbionts of nematodes, which may limit their opportunities for horizontal gene exchange with other bacteria after the establishment of symbiosis.

Our results indicate the bacterial origin of fern IPD113. Surprisingly, AM fungi also possess IPD113 genes. Symbiotic relationships with AM fungi are common in ferns. Among the ferns analyzed in this study, *Adiantum aethiopicum*, *Blechnum occidentale*, *Lygodium flexuosum*, and *Christella dentata* have been reported to exhibit AM colonization (Kumari et al. 2024). The hyphosphere of AM fungi harbors a microbiome distinct from that of bulk soil and the rhizosphere. This microbiome includes *Brevibacillus*, *Cohnella*, *Tahibacter*, *Undibacterium*, *Hymenobacter*, and *Novosphingobium* (Zhang et al. 2022)—all of which have Cry-like two domain proteins and were identified as potential bacterial donors for HGT to AM fungi and ferns (Fig. 6). This is therefore consistent with a scenario of bacteria–AM fungi–fern HGT. Despite the widespread symbiosis of AM fungi with vascular plants, only ferns and bryophytes seem to possess this gene. Understanding the evolutionary significance and underlying mechanisms of this unique distribution in plants will require further investigation.

Ferns appear to have retained IPD113 for its original insecticidal function. However, the physiological roles of proteins containing Cry-like two domains in many bacteria, eukaryotes, and archaea remain unclear. Previous studies have noted that Pra and Prb proteins resemble a beetle protein, which we refer to in this study as the insect Pra+Prb protein (Duchaud et al. 2003; Waterfield et al. 2005). The insect Pra+Prb protein was originally isolated from the hemolymph of the Colorado potato beetle *Leptinotarsa decemlineata* and predicted to function as a juvenile hormone esterase (Vermunt et al. 1997, 1998), although its true function remains unknown. The western corn rootworm *Diabrotica virgifera virgifera* possesses four copies of the gene encoding the three-domain protein, one of which lacks introns (Table S2), suggesting that HGT events have occurred repeatedly, including a relatively recent one. The frequent occurrence of HGT may be attributed not only to close biological interactions but also to the potential contribution of transposable elements. Future studies are needed to elucidate both the functions of proteins containing Cry-like two domains and the genetic mechanisms underlying their broad taxonomic distribution.

## Material and methods

### BLAST searches

Non-redundant protein sequences (nr) were searched using BLASTp (Camacho et al. 2009). The raw output data is shown in Table S1. Taxonomic information was obtained from the NCBI Taxonomy database (Federhen 2012). For downstream analyses, sequences showing ≥95% amino acid identity were removed using CD-HIT (Li and Godzik 2006).

### Structure prediction and alignment

Tertiary structures of full-length BLAST-hit proteins and Cry proteins were predicted using AlphaFold 3 (Abramson et al. 2024) under template-free conditions. Unless otherwise specified, all other parameters followed the default settings of AlphaFold3. For Cry proteins, we selected Cry proteins with quaternary rank 1 from the Bacterial Pesticidal Protein Resource Center (www.bpprc.org). Cry1If1, Cry1Jc1, Cry8La1, and Cry53Aa1 were excluded from the analysis because their amino acid sequences contained unknown residues (X). The top-ranked model among five predictions was used for further analyses. Predicted structures that did not contain complete Cry-like two domains were excluded.

Structural alignments were performed using FoldMason (Gilchrist, Mirdita, and Steinegger 2024). To ensure precise domain trimming, four separate alignments were generated: (1) Cry, (2) BLAST hits obtained using *L. wrightii* IPD113 and *C. faecalis* IPD113-like proteins as queries, (3) BLAST hits obtained using *P. akhurstii* and *V. parahaemolyticus* Prb as queries, and (4) BLAST hits obtained using *P. akhurstii* and *V. parahaemolyticus* Pra as queries. Domain trimming was guided by the reference structures (Cry1Aa1, *L. wrightii* IPD113, *P. akhurstii* Prb, and Pra). For domain I, residues from the N-terminal amino acid forming the first α-helix (α1) to the C-terminal amino acid forming the seventh α-helix (α7) were retained. For domain II, residues from the N-terminal amino acid forming the first β-strand (excluding those included in domain I) to the C-terminal amino acid forming the last β-strand were extracted. The same criteria were applied to Cry domain III and Pra domain. For full-length sequences of the domain I–II region, the linker region between domains I and II was included. After trimming, we evaluated qualities of the trimmed predicted structures using AF3 confidence metrics, pLDDT and predicted aligned error (PAE). For phylogenetic and motif analysis, we used 557 AF3 models with mean pLDDT ≥60 (a per-atom confidence index) to extract amino acid sequences corresponding to domain I, domain II, and the domain I–II region. For structural analysis, we used 498 predicted models with high confidence, defined as an average pLDDT ≥ 80 and PAE ≤ 10 Å in both domains I and II.

### Phylogenetic analysis

Phylogenetic trees were reconstructed using the Graph Splitting method (Figs. 1, 2, S4 and S6) (Matsui and Iwasaki 2020) and the maximum likelihood method (Figs. 5 and S8). For GS trees, we used Graph Splitting v2.4. The Edge Perturbation (EP) values were computed with 100 replicates. For ML trees, the amino acid sequences were aligned using MAFFT v7.526 (Katoh and Standley 2013) and the trees were generated with IQ-TREE2 v2.3.6 (Minh et al. 2020) using the ModelFinder option with 1,000 ultrafast bootstrap and SH-aLRT replicates. The WAG+F+I+R4, pfam+F+R5, and yeast+R5 models were selected for the 2D/3D-mixed, IPD113, and Prb clades, respectively. All trees were visualized using the ggtree R package (Yu et al. 2017).

### Motif analysis

Amino acid motifs were identified using MEME v5.5.7 (Bailey et al. 2015). Preliminary searches suggested that 60 and 20 motifs were sufficient to cover the entire sequences of the two-domain and domain III/Pra/insect Pra-like protein datasets, respectively, with the identified motifs evenly distributed across the alignments. To evaluate whether the identified motifs contained sufficient information and evolutionary relevance, we applied two criteria: (1) average information content ≥ 1.0, and (2) at least two positions with information content ≥ 1.5. Only one motif (motif 40) identified in the domain I–II sequences failed to meet these criteria and was excluded from subsequent analyses. A binary matrix representing the presence or absence of each motif across sequences was constructed and subjected to dimensionality reduction using t-SNE implemented in the Rtsne R package. The perplexity parameter was set to one-third of the number of sequences.

### Structural analysis

Protein structures were visualized using UCSF ChimeraX v1.8 (Pettersen et al. 2004). Pairwise structural alignments were performed in UCSF ChimeraX v1.8 using the MatchMaker command (sequence alignment algorithm, Needleman–Wunsch; Matrix, BLOSUM-62), which provided alignment parameters including sequence score, RMSD (all and pruned pairs), and number of aligned residues.

### Exponential fitting of AA–3Di distance

AA distances were derived from the distance matrices of the ML trees generated using IQ-TREE2. Pairwise distances between MAFFT-aligned 3Di sequences were calculated as *p*-distance, defined as the proportion of differing positions between two sequences after excluding gap positions. Specifically, for each sequence pair !, #, *p*-distance was calculated as:

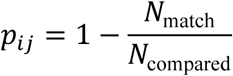

where *N_match_* is the number of aligned positions with identical symbols (excluding gaps), and *N*_compared_ is the total number of positions compared (i.e., sites without gaps in either sequence).

The relationship between sequence and structural similarity is strongly nonlinear, and structural divergence becomes particularly pronounced at low sequence identity; therefore, exponential models have been used to approximate this divergence (Chothia and Lesk 1986). To obtain the bacteria–bacteria regression, we fit an asymptotic exponential model of the form

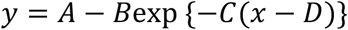

where *x* is the AA distance, y is the 3Di distance, and *A, B, C,D* are free parameters. Nonlinear least-squares fitting was performed using the minpack.lm R package. Outliers beyond ±3 standard deviations of the residuals were removed, and the model was refit using the remaining bacteria–bacteria pairs.

## Acknowledgements

The authors thank Keisuke Shoji (Tokyo University of Agriculture and Technology) for fruitful discussion. This research was partially supported by the Japan Society for the Promotion of Science (JSPS) KAKENHI (25K22485).

## Author Contributions

H.E. designed research, performed analysis and wrote the manuscript. H.K. contributed to data interpretation and critically reviewed the manuscript.

## Conflict of interest

The authors declare no conflict of interest.

## Data availability

All data generated and used for the analyses are deposited in the CBS Data Sharing Platform (URL and password are provided for reviewers and it will be made public upon acceptance)

## Code availability

Custom code used in the analysis is deposited in the CBS Data Sharing Platform (URL and password are provided for reviewers and it will be made public upon acceptance). Protein structure modeling and structural alignment were performed using AlphaFold 3 (https://github.com/google-deepmind/alphafold3) and FoldMason (https://github.com/steineggerlab/foldmason). Phylogenetic analysis was conducted using the Graph Splitting method (https://github.com/MotomuMatsui/gs) and IQ-TREE2 (https://github.com/iqtree/iqtree2). Amino acid motif analysis was performed using MEME (https://github.com/cinquin/MEME). Sequence alignment and processing were performed using MAFFT (https://github.com/GSLBiotech/mafft) and CD-HIT (https://github.com/weizhongli/cdhit). For plotting and data processing, we used Python v.3.10.18 (https://www.python.org/), R v.4.4.1 (https://www.r-project.org/), and R studio v2024.12.1+563.

---

**Fig S1.**
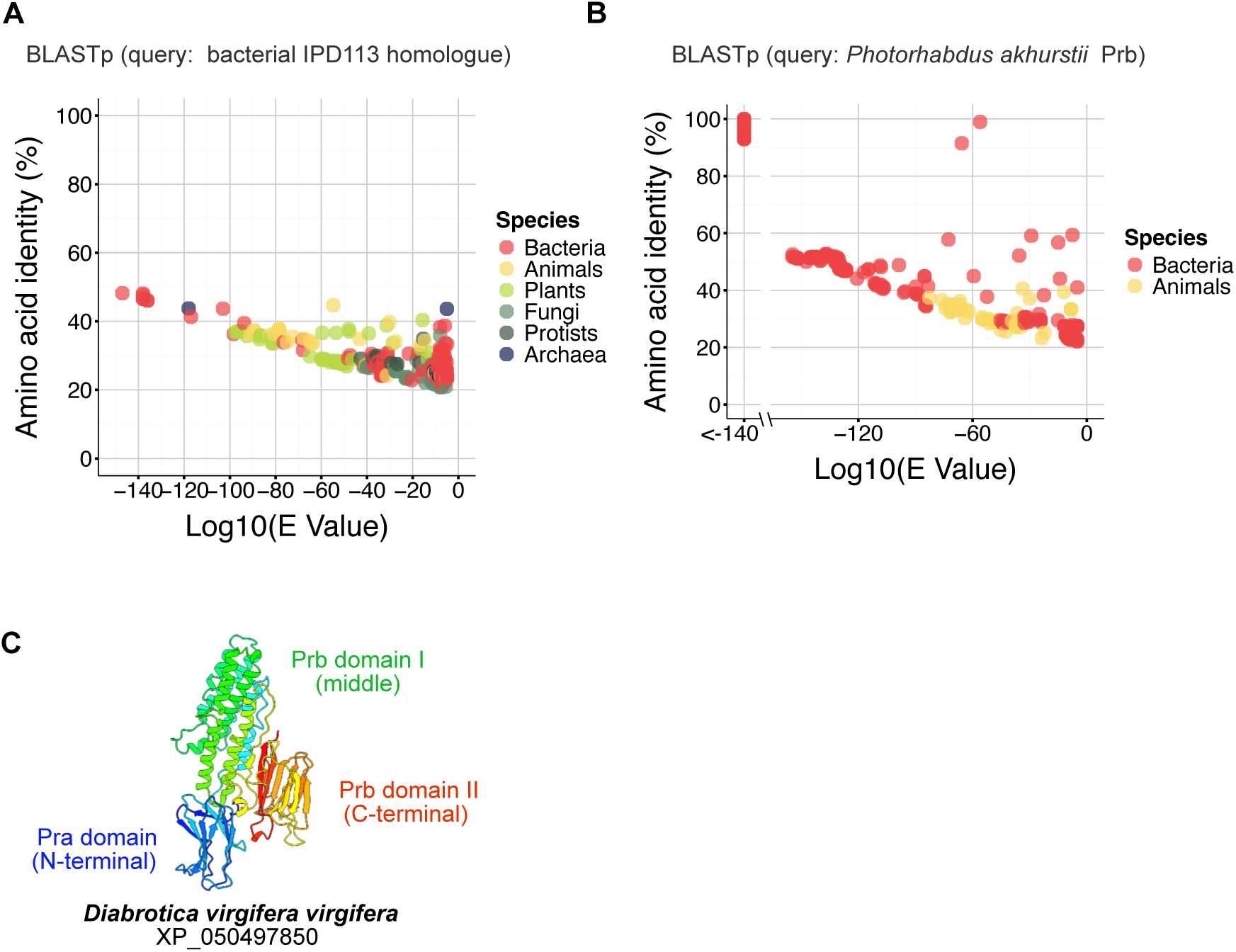
Putative IPD113 and Prb homologues identified by BLASTp. (A, B) Scatter plots of amino acid identity versus E-value from BLASTp searches. *Cohnella faecalis* IPD113-like protein (A) and *Photorhabdus akhurstii* Prb protein (B) were used as queries. Dots indicate putative homologues (E-value < 1e−5). Raw BLASTp results are provided in Table S1. (C) AF3-predicted structures of a putative Prb homologue.

**Fig S2.**
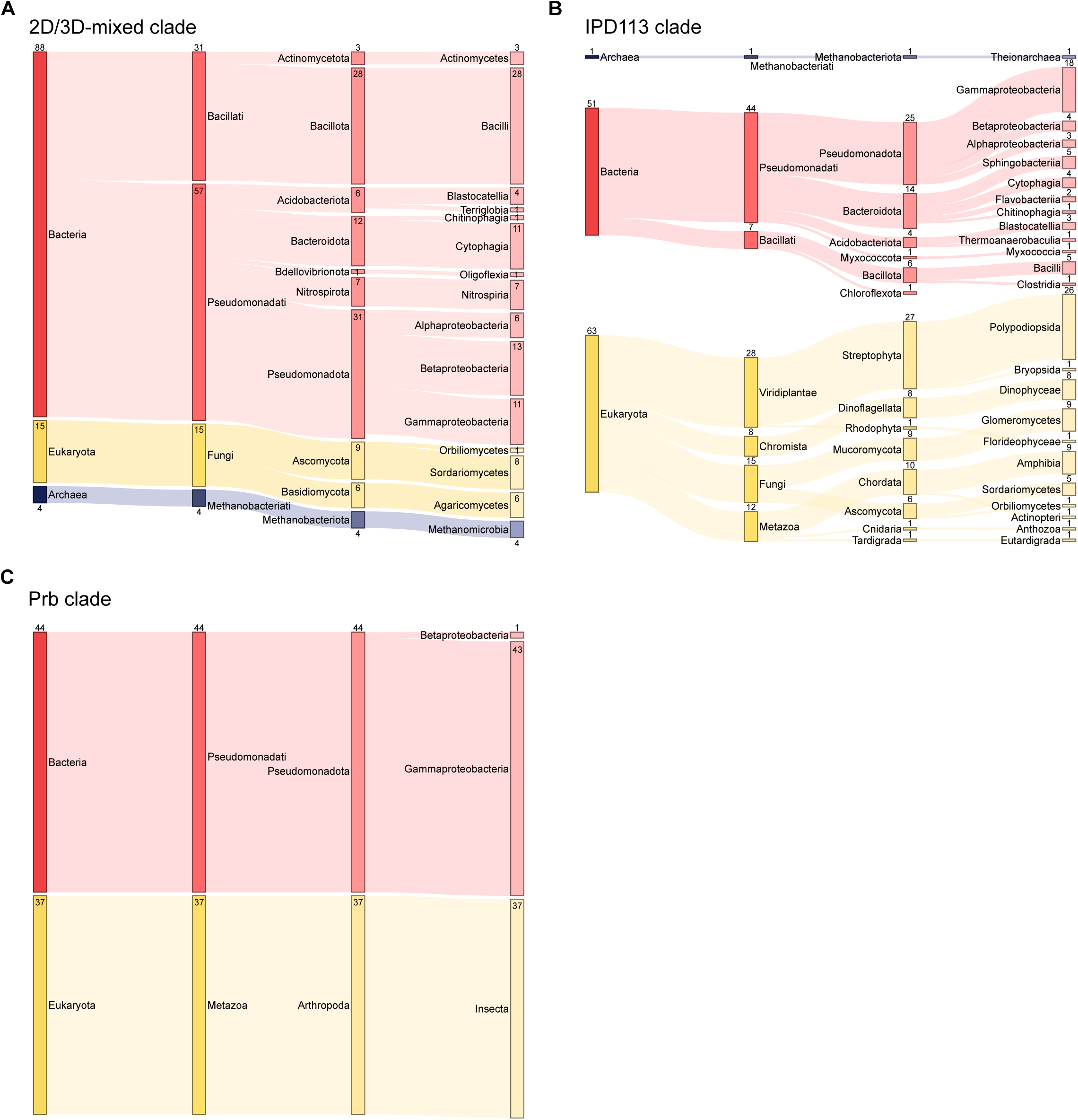
Sankey plots showing the taxonomy of proteins in the 2D/3D-mixed (A), IPD113 (B), and Prb (C) clades. Full taxonomic assignments derived from NCBI are listed in Table S2.

**Fig S3.**
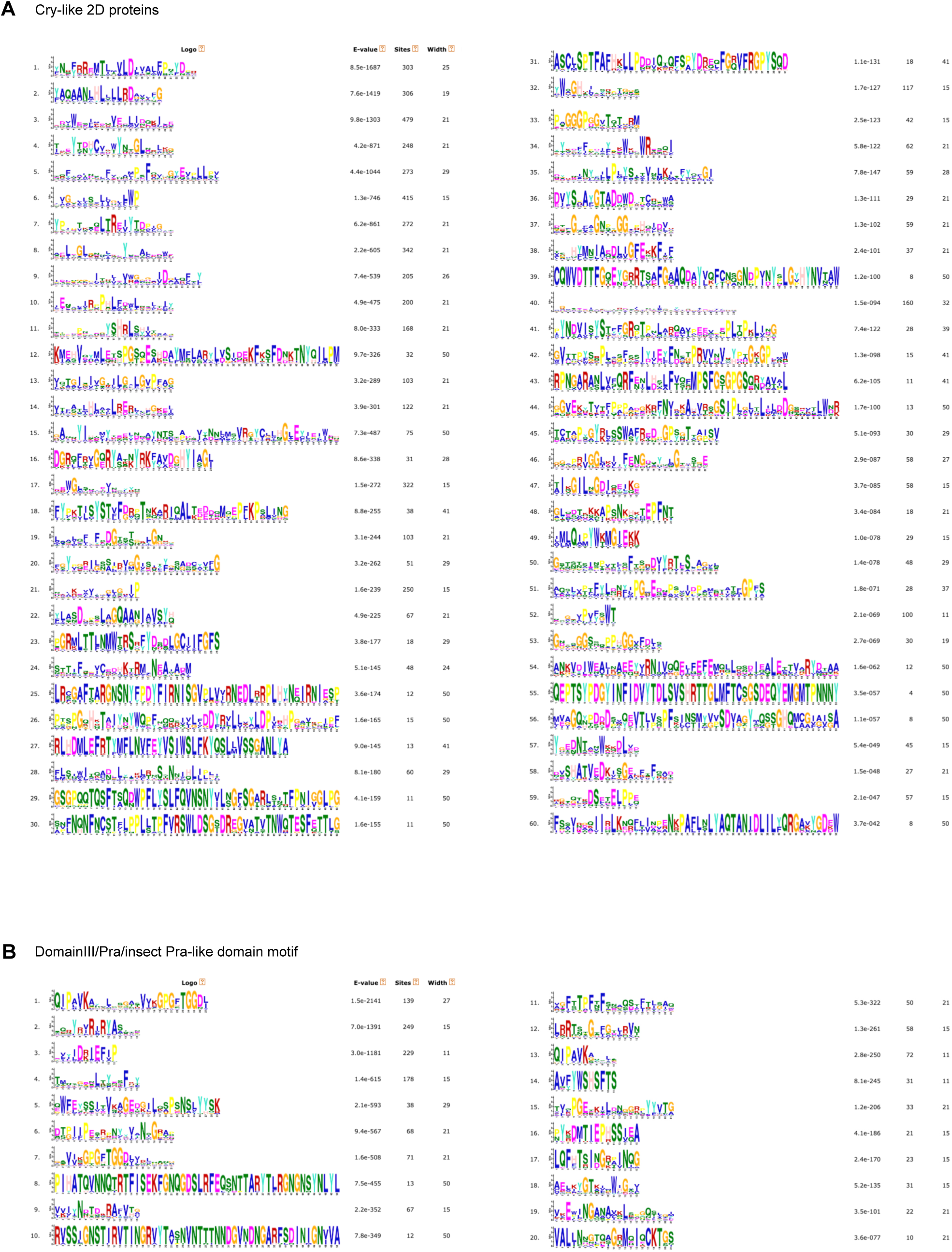
Amino acid motifs identified by MEME. Amino acid motifs of (A) Cry-like 2D proteins and (B) domain III, Pra, insect Pra-like domain proteins.

**Fig S4.**
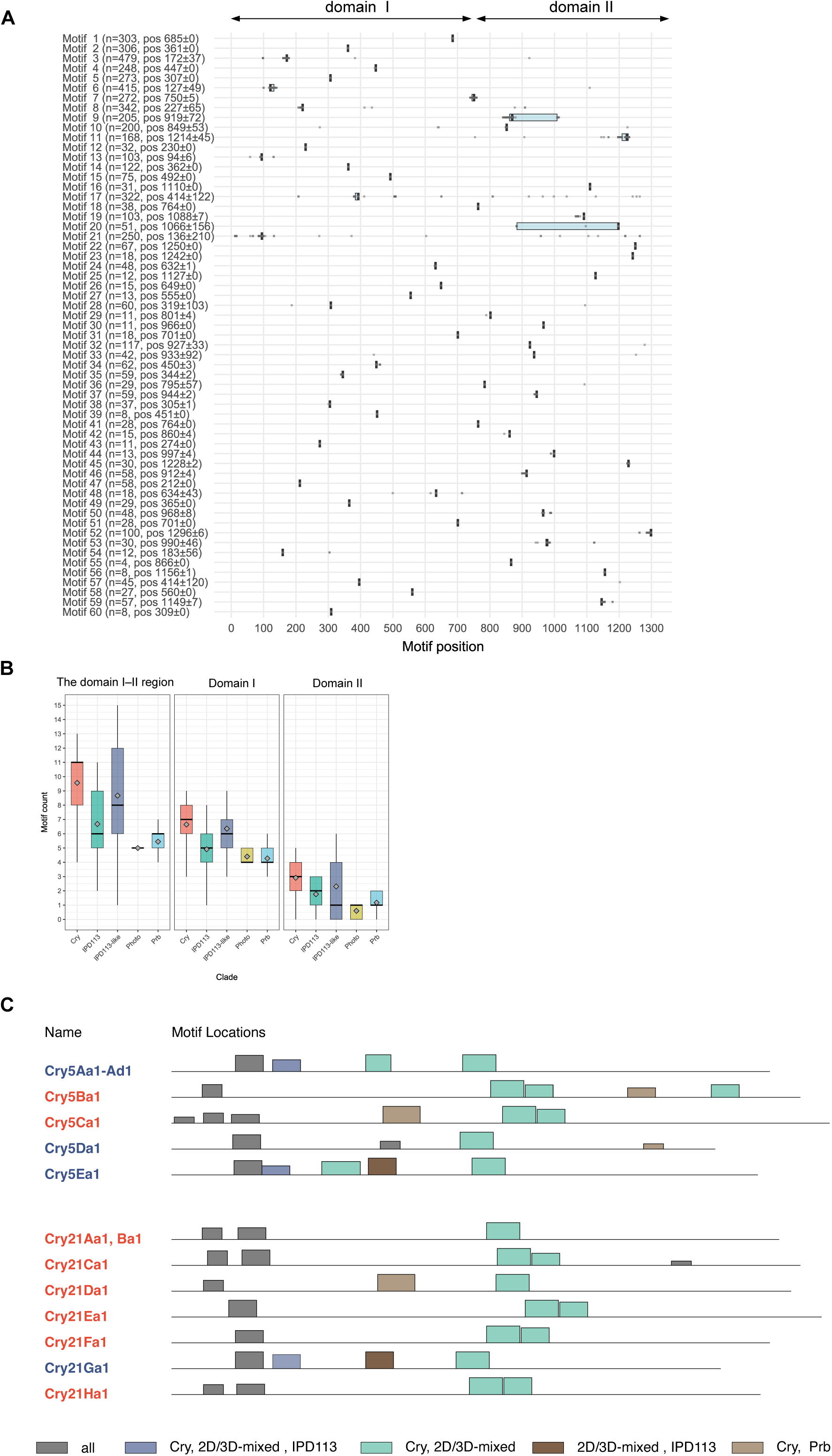
Positions and counts of amino acid motifs. (A) Distribution of motif positions mapped onto the structural alignment generated by FoldMason. Each row corresponds to one detected motif. Labels show the number of occurrences (*n*) and the mean starting position of the motif ± standard deviation. Motif 40 was excluded from the analysis because it did not meet two information-content criteria (see Methods). Light blue boxplots show the median (vertical black line), first and third quartiles (box), and whiskers extending to 1.5× the interquartile range. Gray dots represent individual occurrences of each motif. For most motifs, the interquartile range is extremely small, making the light blue boxes barely visible. (B) Counts of motifs in domain I, domain II, and the domain I–II region among members of each clade. Boxplots show the median (horizontal line), first and third quartiles (box), and whiskers extending to 1.5× the interquartile range (outliers hidden); diamonds indicate the mean. (C) Motif locations in Cry5 and Cry21 proteins. Each rectangle represents a single motif, with colors indicating the combination of clades in which the motif is shared.

**Fig S5.**
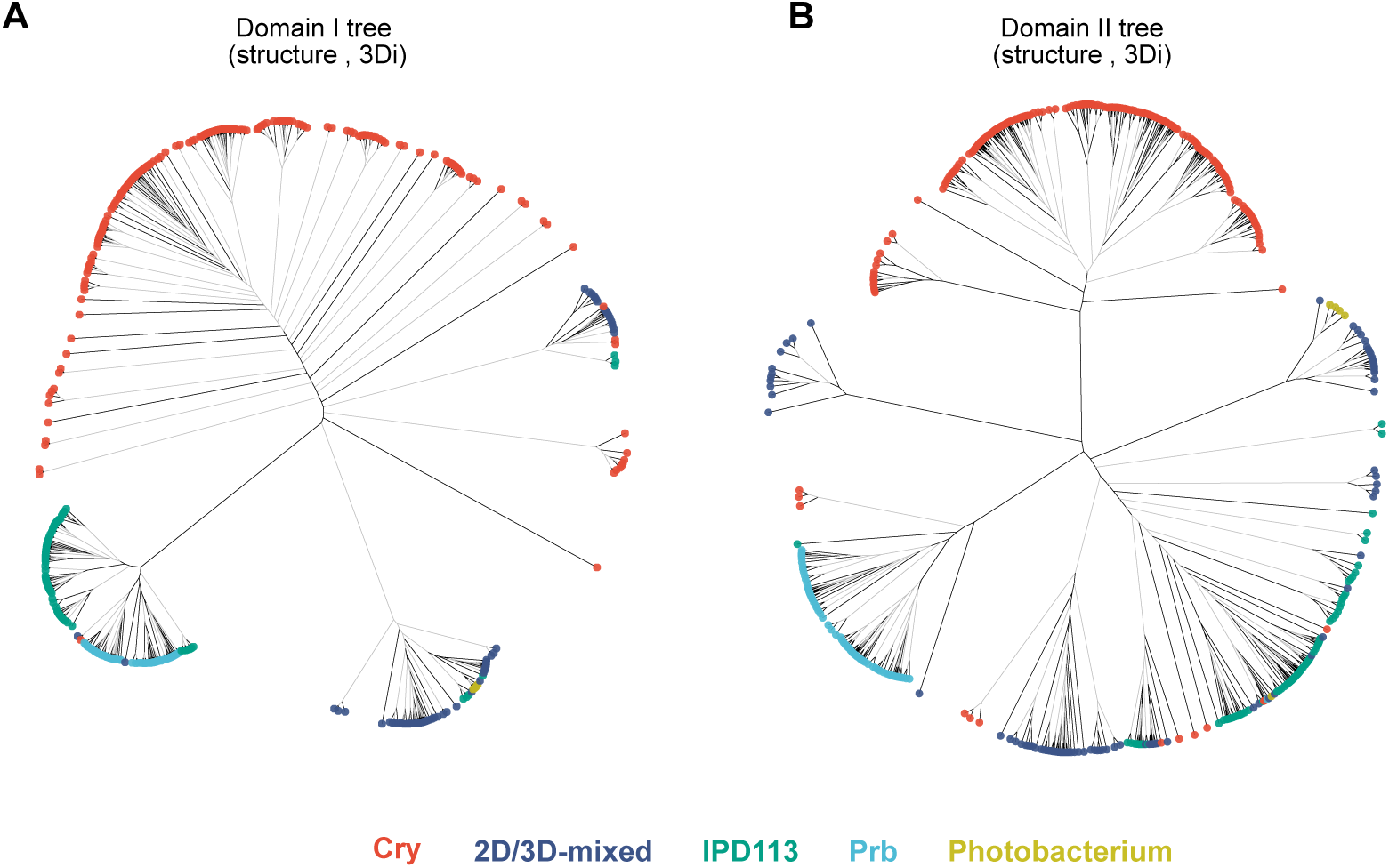
Domain-level GS trees based on 3Di sequences. (A) Domain I tree. (B) Domain II tree. Black and gray branches indicate high and low reliability, supported by edge perturbation (EP) values ≥ 0.9 and < 0.9, respectively. Node circle colors correspond to the clades defined in the two-domain GS tree shown in Fig. 1C.

**Fig S6.**
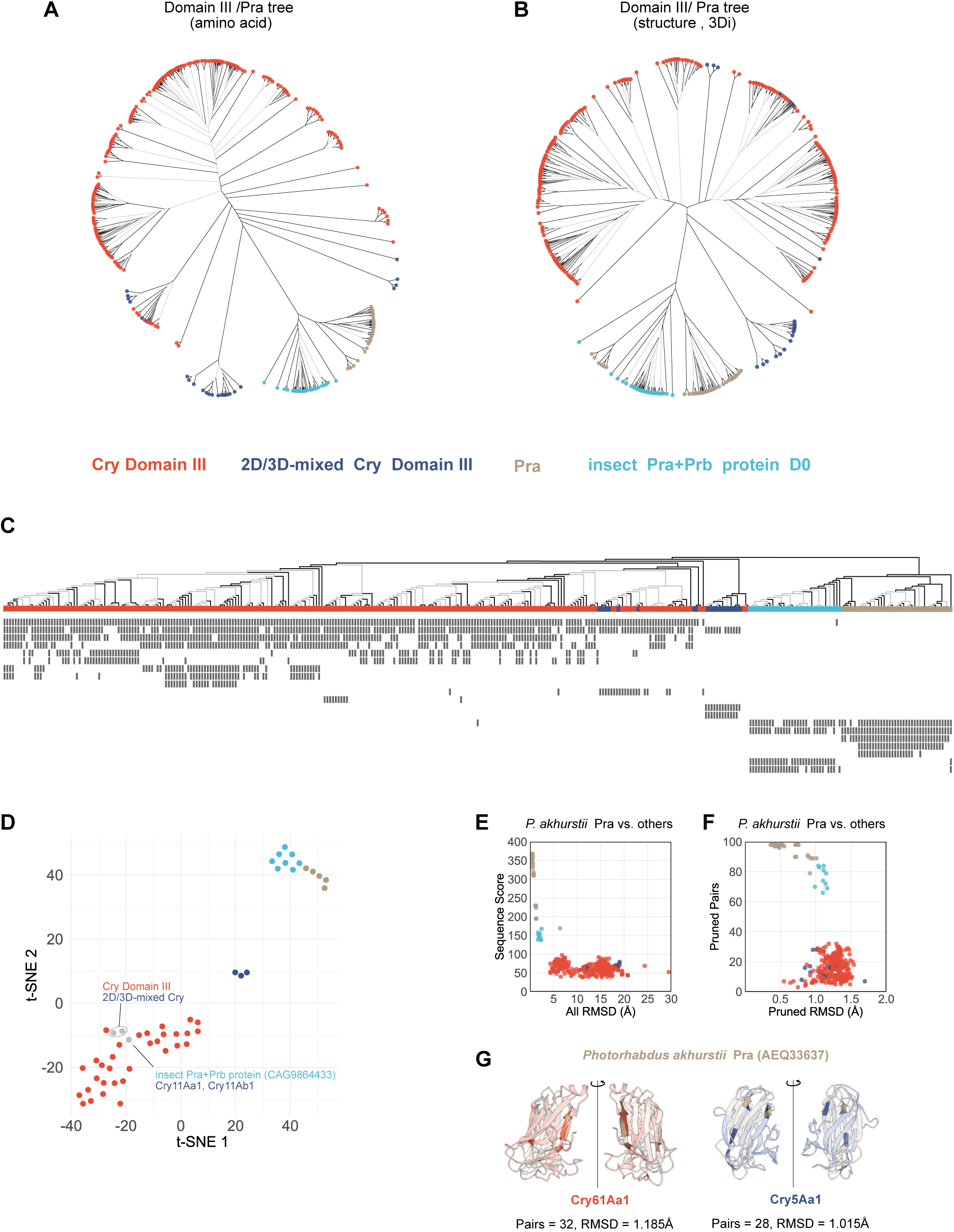
Analyses of Cry domain III, Pra, and insect Pra-like domains. (A, B) GS trees of Cry domain III, Pra, and insect Pra-like domains based on amino acid (A) and 3Di (B) sequences. Black and gray branches indicate high and low reliability, supported by edge perturbation (EP) values ≥ 0.9 and < 0.9, respectively. Node circle colors correspond to the clades. (C) Raster plot of amino acid motif patterns. Twenty motifs were mapped onto the GS tree shown in (A). (D) t-SNE visualization of motif patterns. The motif presence/absence matrix from (A) was dimensionally reduced. Sequences without motifs (Cry11Ba1, Cry18Aa, Cry18Ba1, Cry18Ca1) were excluded from this analysis. Dot colors correspond to the same clade color scheme as in (A) and (B). (E–G) Structural comparisons of *P. akhurstii* Pra with other members shown in (A). Colors correspond to the same clade color scheme as in (A). (E) Sequence similarity scores and whole-domain RMSD values obtained from pairwise alignments and structural superpositions. (F) Pruned residue pairs and their pruned RMSD values derived from structural superpositions. (G) Representative structural superpositions with proteins from different clades. Pruned regions are highlighted, and non-pruned regions are shown with increased transparency.

**Fig S7.**
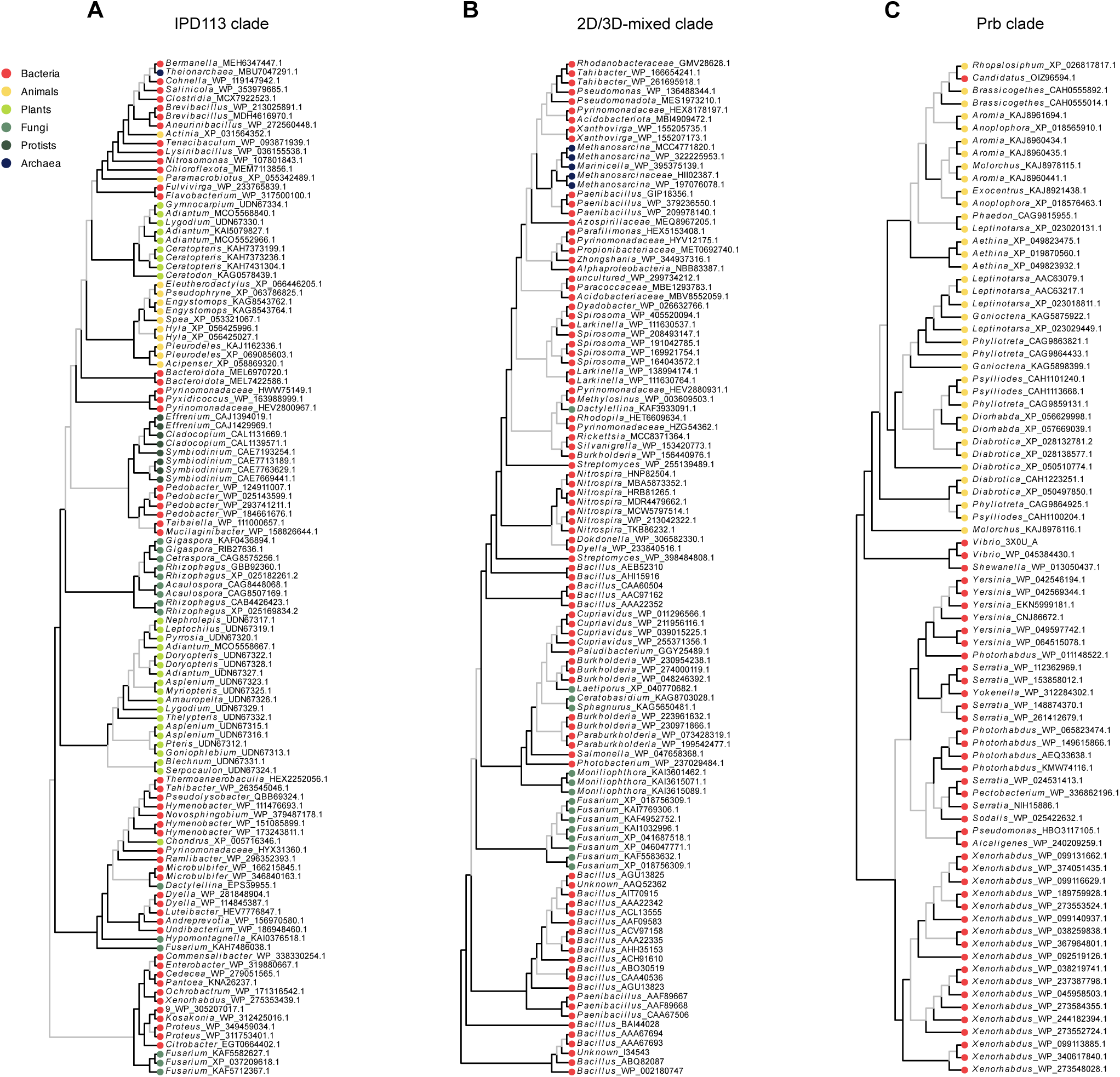
The two-domain GS tree colored by taxonomy. The two-domain GS tree shown in Fig. 1C is colored by taxonomy for the IPD113 (A), 2D/3D-mixed (B), and Prb (C) clades. Black and gray branches indicate high and low reliability, supported by edge perturbation (EP) values ≥ 0.9 and < 0.9, respectively.

**Fig S8.**
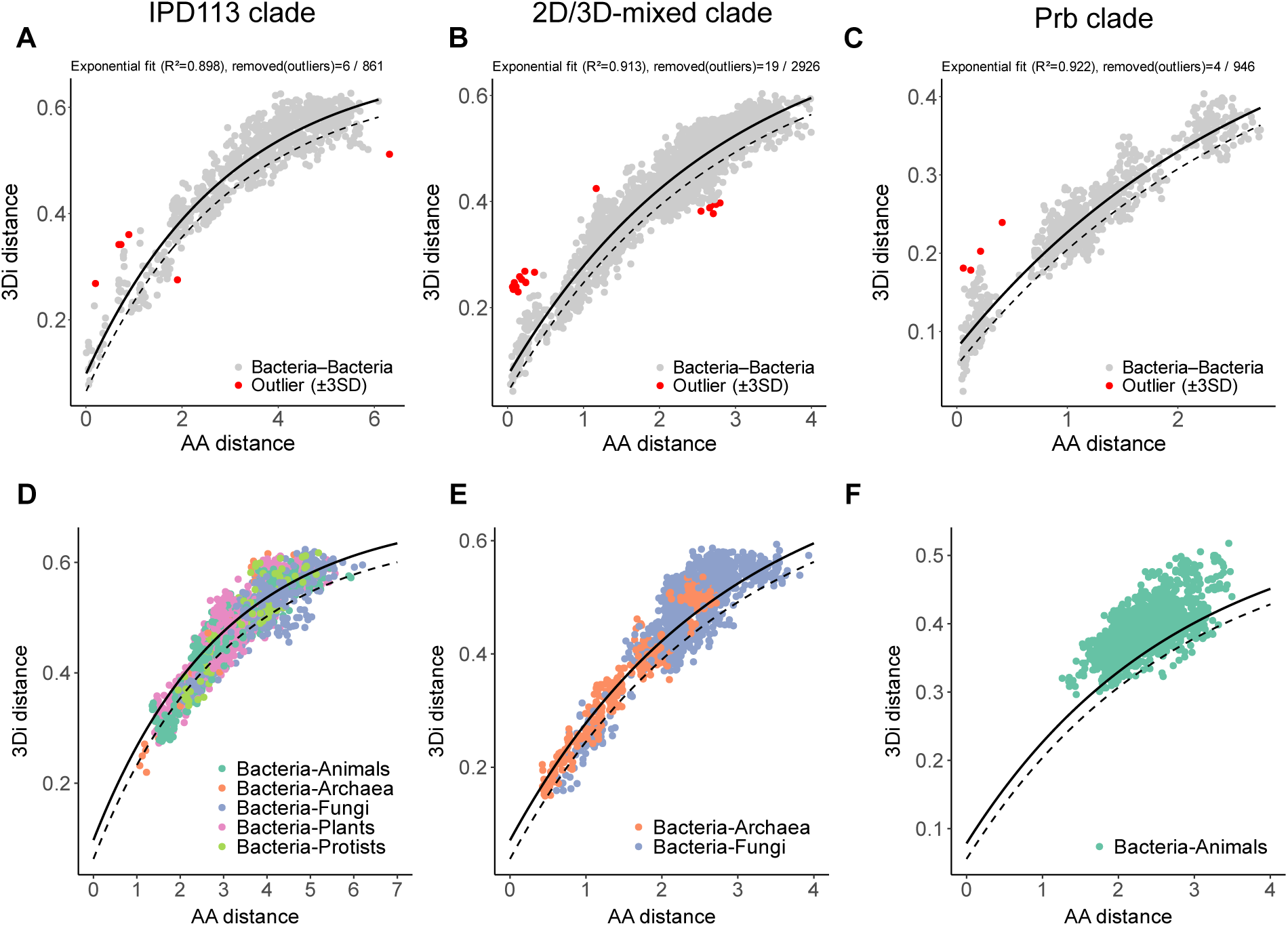
Distributions of AA and 3Di distances. (A–C) Distributions of amino acid (AA) and 3Di distances among bacteria–bacteria pairs in the IPD113 (A), 2D/3D-mixed (B), and Prb (C) clades. AA distances were derived from the distance matrices of the ML trees shown in Figs. 5 and S9A and C. 3Di distances were calculated as pairwise distances (p-distances) based on 3Di alignments. Black solid lines indicate the exponential fit; dotted lines indicate fit –1 SD. Red dots represent outliers. (D–F) Distributions of AA and 3Di distances among all pairs in the IPD113 (D), 2D/3D-mixed (E), and Prb (F) clades. Black solid lines indicate the exponential fit; dotted lines indicate fit –1 SD. Dots representing pairs between bacteria and non-bacterial taxa are color-coded.

**Fig S9.**
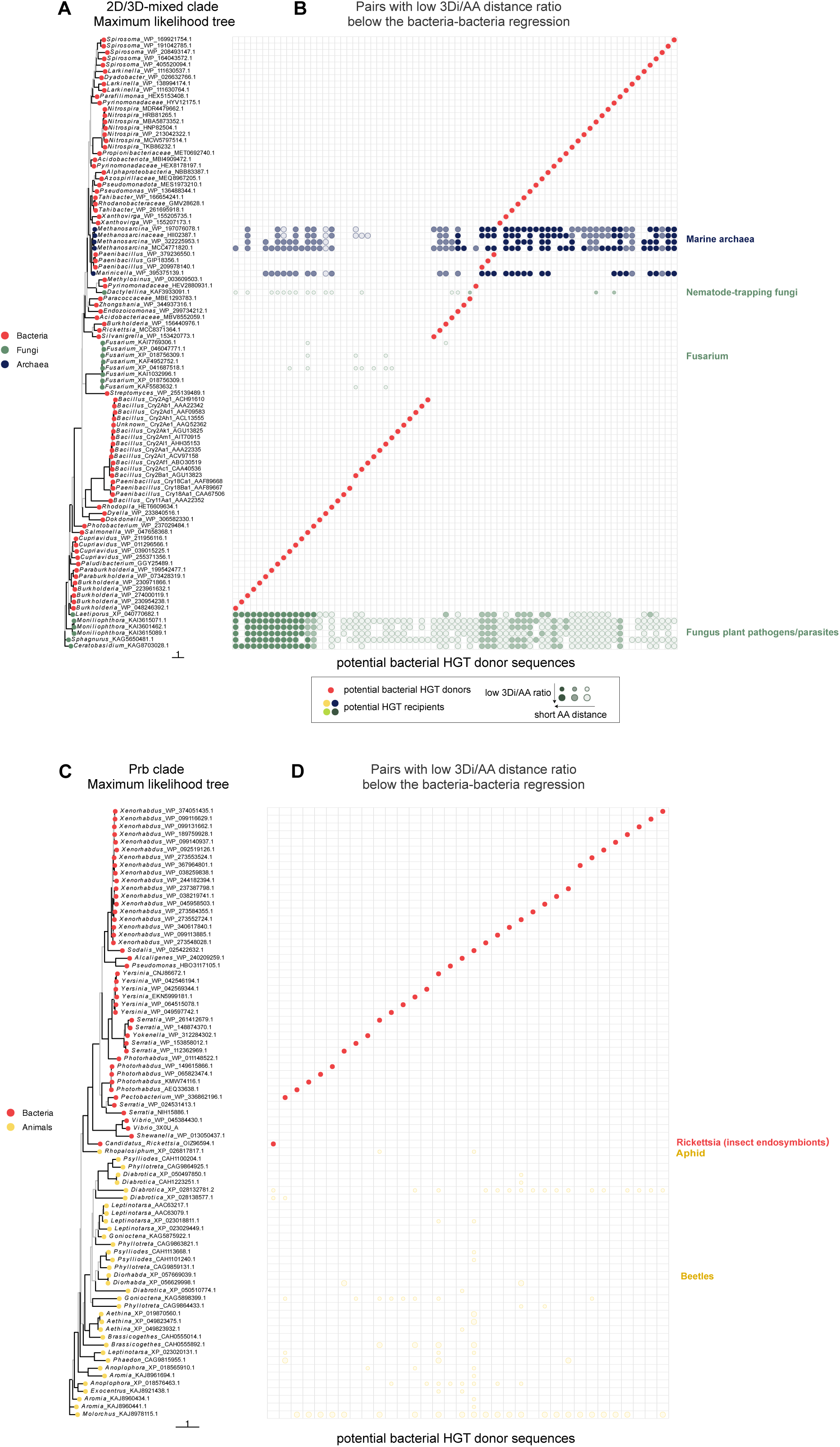
Potential HGT pairs in the 2D/3D-mixed and Prb clades. (A, C) Maximum likelihood tree of the 2D/3D-mixed and Prb clades. Black branches indicate high reliability (Ultrafast Bootstrap ≥ 95% and SH-aLRT ≥ 80%); other branches are shown in gray. Node circle colors indicate taxonomy. The scale bar represents the number of substitutions per site. (B, D) Pairs with a low 3Di/AA distance ratio, falling below the bacteria–bacteria regression. The vertical position of each circle corresponds to the sequence labels in the phylogenetic tree shown in (A). Each column corresponds to a red-highlighted bacterial sequence (potential HGT donor). Proteins plotted in the same column have a 3Di/AA distance ratio that falls below the bacteria–bacteria regression in Fig. S8, indicating potential HGT recipients. The fill opacity of each circle reflects its amino acid (AA) distance relative to the mean of bacteria–bacteria pairs: circles with AA distances above the mean are outlined with high transparency, those between the mean and −1 SD are semi-transparent, and those below −1 SD are shown as fully opaque. Large circles indicate proteins below −1 SD from the regression, while small circles indicate those between 0 and −1 SD. In particular, large opaque circles are more likely to represent candidate HGT pairs involving bacterial sequences.

**Table S1 | Raw BLASTp results**

**Table S2 | List of proteins containing Cry-like two domains used in this study.**

## Notes

### Competing Interest Statement

The authors have declared no competing interest.

### Summary of Updates

Line numbers added; Data availability updated.

